# PRESYNAPTIC MUSCARINIC-NICOTINIC MODULATION of GABAergic INPUTS to NEURONS in the REM SLEEP EXECUTIVE AREA of the ROSTRAL PONS

**DOI:** 10.1101/2025.06.09.658683

**Authors:** Esteban Pino, Héctor Kunizawa, Michel Borde

## Abstract

A large body of evidence indicates that cholinergic neurons in the brainstem tegmental nuclei play a pivotal role in controlling neurons of the reticular nucleus pontis oralis (PnO), an area considered executive for rapid eye movement (REM) sleep onset and maintenance. More recent data indicate that the PnO is also under GABAergic control -which gates REM sleep- thereby highlighting the importance of interactions between cholinergic and GABAergic processes as a key mechanism in REM sleep regulation. Here we employed a rat mesopontine slice preparation to investigate, *in vitro*, the modulation of GABAergic inputs to PnO neurons by cholinergic agents. Carbachol, a mixed muscarinic-nicotinic cholinergic agonist, provoked either depression or facilitation of single monosynaptic GABAergic IPSCs evoked by extracellular stimulation of the region of brainstem tegmental nuclei. Both effects were presynaptic in origin and are likely attributable to the activation of distinct presynaptic cholinergic receptors as depression was replicated by muscarine (presynaptic inhibition) and facilitation by nicotine (presynaptic facilitation). Furthermore, IPSCs evoked by stimulation patterns mimicking physiological activity (short trains at 15 Hz) were also affected by cholinergic agonists. Muscarine caused presynaptic inhibition accompanied by frequency facilitation and nicotine promoted the opposite effect. Notably, both agonists reduced the total inhibitory charge transferred to postsynaptic neurons during the stimulus train. These findings suggest an additional mechanism by which cholinergic modulation can relieve GABAergic inhibition of PnO neurons under physiological conditions, serving as a local component of the reciprocal inhibitory interactions between sleep-regulating networks in REM sleep brainstem executive areas.

**NEW & NOTEWORTHY:** Despite decades of research into brainstem sleep mechanisms, the GABAergic-cholinergic interactions involved in gating and ultradian regulation of REM sleep remain poorly understood. We provide evidence for the functional expression of presynaptic muscarinic and nicotinic cholinergic receptors at GABAergic terminals within the pontine reticular nucleus (PnO), a region central to REM sleep onset and maintenance. Our results indicate that cholinergic signaling might relieve GABAergic inhibition of PnO neurons. As a component of a mutual inhibitory mechanism between REM sleep controlling systems, this interaction may sharpen transitions into REM sleep.

## INTRODUCTION

Mesopontine neural networks are critically involved in organizing various sleep-wake states, such as rapid eye movement (REM) sleep and wakefulness (W) (Brown et al., 2012; Jones, 2020). A wealth of experimental evidence supports the role of various neurotransmitters and neuromodulators in controlling the occurrence and characteristics of these states (Xi et al., 1999; 2004; Fuller et al., 2007; Luppi et al., 2006; Arrigoni et al., 2016; Scammell et al., 2017; Jones, 2020; Wang et al., 2021). Electrophysiological, pharmacological, immunohistochemical and behavioral data indicate that cholinergic and GABAergic processes converge at these pivotal mesopontine areas and interact locally to facilitate either REM sleep or W. Moreover, the interplay between GABAergic and cholinergic systems may play a significant role in establishing the ultradian rhythmicity of sleep (Xi et al., 1999; 2004; Standford et al., 2003; Liang and Marks; 2009; Brown et al., 2008; Marks and Sinton, 2010; Reinoso-Suárez et al., 2010; Vanini et al., 2011). Nonetheless, the cellular and synaptic mechanisms underlying these interactions remain poorly understood.

Although the precise role of brainstem cholinergic transmission in regulating REM sleep remains debated, there is broad consensus that cholinergic neuromodulation contributes to its control (for reviews, see Marks & Sinton, 2010; Brown et al., 2012; Jones, 2020). Diverse experimental approaches -including optogenetics, pharmacogenetics, genetic manipulations for imaging and selective deletions, as well as virus-mediated circuit tracing-have allowed to identify a conserved brainstem network located in the rostral pons that is critically involved in REM sleep regulation. This network comprises a distinct group of cholinergic neurons situated in the brainstem tegmental nuclei -namely, the laterodorsal (LDT) and pedunculopontine tegmental (PPT) nuclei-and their projections to glutamatergic neurons in the reticular nucleus pontis oralis (PnO), which is considered pivotal for REM sleep onset and maintenance (Reinoso-Suárez et al., 1994; Garzón et al., 1998). More recent studies have demonstrated that REM sleep–regulating circuits also include specific glutamatergic neurons within the pontine tegmentum, such as the peri-locus coeruleus α (peri-LCα) region in cats and the subcoeruleus (subLC) or sublaterodorsal nucleus (SLD) in rodents (Arrigoni et al., 2016; Saper and Fuller; 2017). Historically, cholinergic inputs to these brainstem regions were seen as essential for initiating REM sleep (Brown et al., 2012; Weber and Dan, 2016). However, more recent findings suggest that brainstem cholinergic neuronal groups primarily act as modulators rather than as generators of REM sleep, with glutamatergic and GABAergic transmissions being the main players in REM sleep generation and maintenance (Luppi et al., 2006; Scammell et al., 2017; Niwa et al., 2018; Arrigoni & Fuller, 2019; Wang et al., 2021).

GABA has long been regarded as a sleep-promoting neurotransmitter. However, extensive research led to the incorporation of GABAergic transmission into models that explain not only the occurrence of REM sleep episodes but also the sequence of different wakefulness–sleep stages (for a review see Wang et al., 2021). The primary source of GABAergic inputs to the REM sleep– executive areas in the pons has been identified as populations of neurons located in the ventrolateral periaqueductal gray (vlPAG) and the lateral pontine tegmentum (LPT) (Boissard et al., 2003; Lu et al., 2006; Sapin et al., 2009; Weber et al., 2018; Luppi et al., 2024). Additional inputs may also arise from GABAergic cells in the PnO and non-cholinergic neurons within the LDT-PPT complex (Semba, 1993; Boissard et al., 2003; Lu et al., 2006; Sapin et al., 2009; Pino et al., 2017). Seminal behavioral evidence for a GABAergic pontine mechanism controlling W and REM sleep comes from experiments in unanesthetized chronic cats. These studies demonstrated that pontine microinjections of GABA or its agonists induced prolonged wakefulness, whereas administration of a GABA-A receptor antagonist resulted in long-lasting REM sleep episodes (Xi et al., 1999). Subsequent rodent studies further supported a critical role for pontine GABAergic mechanisms in REM sleep control (Boissard et al., 2002; Pollock and Mistlberger, 2003; Sanford et al., 2003; Fenik and Kubin, 2009). Current evidence supports the proposal by Xi et al. (2001) that a pontine gating mechanism promotes wakefulness when the tonic activation of GABAergic neurons is high and permits REM sleep when GABAergic tone is low (Sanford et al., 2003; see reviews in Arrigoni et al., 2016 and Wang et al., 2021). Furthermore, this gating mechanism likely involves a mutual inhibitory interaction between W-promoting GABAergic and REM sleep-promoting cholinergic processes operating in REM sleep executive brainstem areas (Xi et al., 2004; Vazquez and Baghdoyan, 2004; Fuller et al., 2007; Fenik and Kubin, 2009; Vanini et al., 2011, 2012; Weber et al., 2018).

Although postsynaptic opposing effects of GABA and cholinomimetic drugs on PnO neurons have been reported (Nuñez et al., 1997; 1998; Brown et al., 2006; Heister et al., 2009; Weng et al., 2014), most research on GABAergic and cholinergic interactions in REM sleep-executive brainstem areas suggests that these interactions primarily occur at the presynaptic level. *In vivo*, the suppression of REM sleep by local application of GABA at the PnO relies on the inhibition of acetylcholine (ACh) release from cholinergic terminals (Vazquez and Baghdoyan, 2004; Marks et al., 2008; Flint et al., 2010). Interestingly, in support of this model, GABA-A receptors have been identified on cholinergic varicose axons in the PnO (Liang and Marks, 2014). In contrast, presynaptic regulation of GABAergic transmission at the PnO mediated by acetylcholine (ACh) or its agonists has been less extensively investigated, and the *in vitro* findings remain inconclusive (Heister et al., 2009).

Based on the concept of a local, mutual inhibitory interaction between GABAergic and cholinergic neurotransmission in the PnO, we thus hypothesized that cholinergic inputs exert inhibitory control over GABAergic inputs to PnO neurons. We tested this hypothesis, in an *in vitro* rat brainstem slice preparation containing the mesopontine structures critically involved in the control of REM sleep (Pino et al., 2017). Our data revealed the presence of presynaptic muscarinic (mAChRs) and nicotinic (nAChRs) acetylcholine receptors on GABAergic terminals contacting PnO neurons. We observed that the activation of mAChRs and nAChRs exerted opposite effects on GABA release probability, respectively. Strikingly, under conditions of repetitive presynaptic stimulation, activation of either receptor type ultimately reduced the overall inhibitory drive. These results support and extend the concept of a local, mutual inhibitory interaction between GABAergic and cholinergic processes in the REM sleep-executive brainstem areas.

## MATERIALS AND METHODS

Wistar rat pups were obtained from URBE (Unidad de Reactivos Biológicos de Experimentación, School of Medicine, Universidad de la República, Uruguay). All experimental procedures adhered to ASAP/ABS Guidelines for the Use of Animals in Research and were approved by the Uruguayan Committee on Animal Research (Comisión Honoraria de Experimentación Animal, CHEA) under Protocol Number 070153-000741-15.

### Mesopontine slice preparation

Mesopontine slices (∼350 µm thick) containing the LDT/PPT complex, the PnO, and the trigeminal motor nucleus (MV) were obtained from 7- to 15-day-old Wistar rats using previously described methods (see Pino et al., 2017). Briefly, the animals were decapitated, a craniotomy was performed, and the brainstem was excised by transection following cerebellum removal. Throughout the procedure, the preparation was continuously bathed in modified artificial cerebrospinal fluid (ACSF) at ∼4°C, containing (in mM): 213 C_₁₂_H_₂₂_O_₁₁_, 2.7 KCl, 1.25 H_₂_KPO_₄_, 2 MgSO_₄_•7H_₂_O, 26 NaHCO_₃_, 10 C_₆_H_₁₂_O_₆_, and 2 CaCl_₂_. Transverse mesopontine slices were sectioned at an inclination of approximately 4–6 degrees relative to the coronal plane using a Vibratome (Vibratome 1000plus sectioning system). Consequently, the dorsal portion of the slice was more anterior (∼-8.72 mm from bregma) than the ventral aspects (∼-8.30 mm; see Paxinos & Watson, 1998). Slices were stored in a holding chamber containing a 50:50 mixture of modified ACSF and standard ACSF (composition in mM: 124 NaCl, 2.7 KCl, 1.25 H_₂_KPO_₄_, 2 MgSO_₄_•7H_₂_O, 26 NaHCO_₃_, 10 C_₆_H_₁₂_O_₆_, and 2 CaCl_₂_) at room temperature (21–24°C). The ACSF pH was maintained at 7.42 by gentle bubbling with carbogen (95% O_₂_, 5% CO_₂_), and the mixture was gradually replaced (∼1.5 h) with 100% standard ACSF. Slices were then transferred to a recording chamber mounted on an upright microscope stage (Nikon Eclipse FN1), equipped with infrared (IR) and differential interference contrast (DIC) imaging devices, as well as ×4 (air) and ×40 (water immersion) objectives. During recording, slices were superfused at a flow rate of ∼2–3 ml/min with standard ACSF at room temperature (21–24°C).

### Recording and stimulation

Whole-cell patch recordings (continuous single-electrode voltage clamp mode) were obtained from visually identified PnO neurons using IR-DIC optics. The recording area was located approximately 250 µm medial to the trigeminal motor nucleus, 250 µm lateral from the midline, and 1.25 mm dorsal to the slice’s ventral boundary (See Fig.1 in Pino et al., 2017). Patch pipettes (3-5 MΩ) were pulled from borosilicate glass (BF150-86-10, Sutter Instrument Co.) using a programmable puller (P-87 Sutter Instruments Co.) and filled with ‘patch solution’ containing (in mM): 150 KCl, 4.6 MgCl_2_, 10 HEPES, 1 EGTA, 0.1 CaCl_2_, 4 ATP-Na, 0.3 GTP-Na. The pipette holder was attached to the headstage of an Axoclamp 2B amplifier (Molecular Devices, San José, CA, USA) and positioned under visual guidance using a hydraulic micromanipulator (Narishige MHW-13). Signals were low-pass filtered (5 KHz, FL-4 DAGAN Corporation), digitized (40 KHz) using a Digidata 1322A analog-to-digital DAQ device (Molecular Devices, San José, CA, USA) and recorded to a PC using pClamp 8.0 software (Molecular Devices, San José, CA, USA).

**Figure 1.**
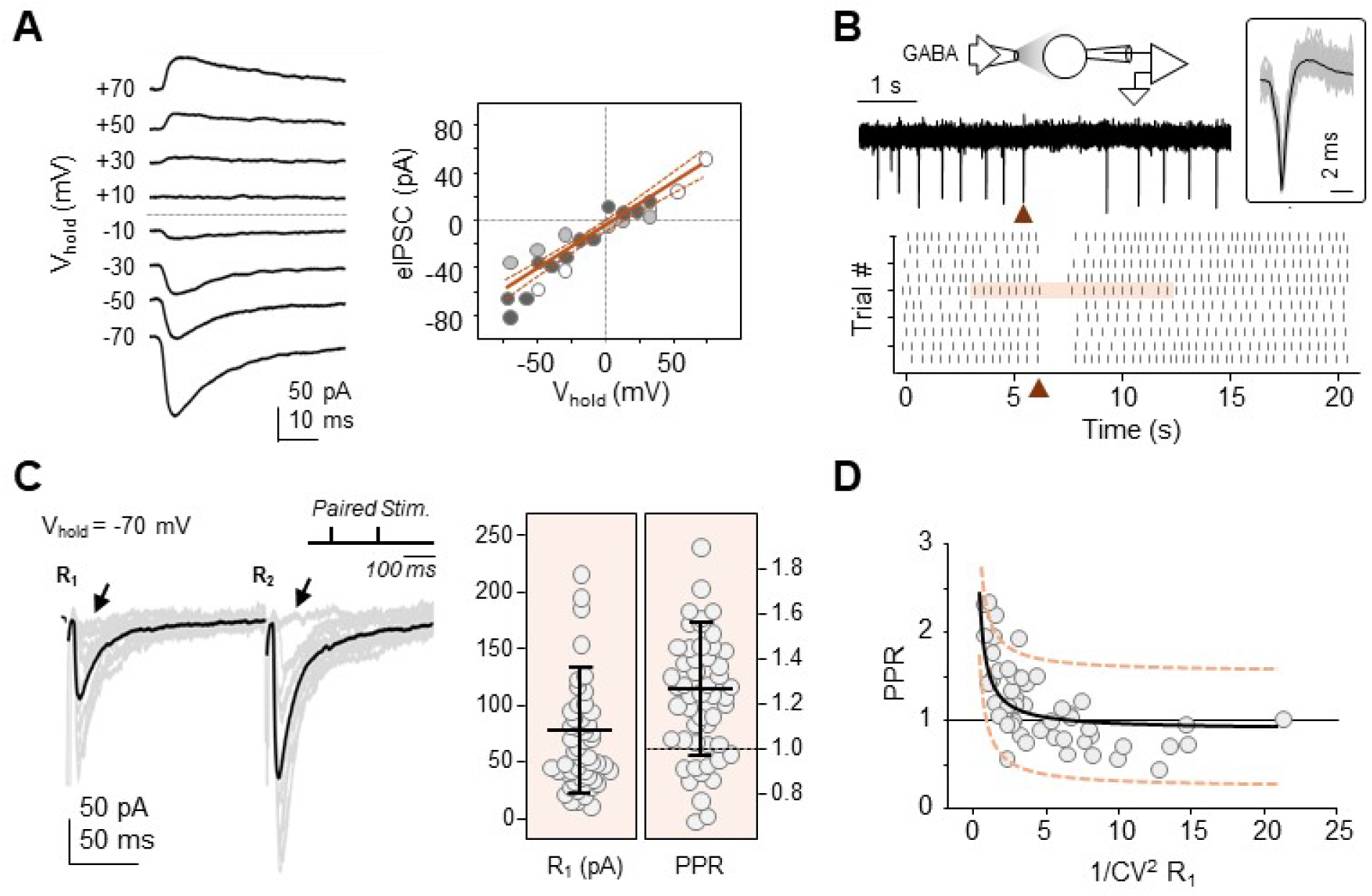
Main functional features of evoked inhibitory postsynaptic currents (eIPSCs). **A**. **Left:** Representative eIPSCs recorded at holding potentials (V_hold_) ranging from -70 to +70 mV, in 20 mV increments. **Right:** current-voltage relationship (N=3) with the best-fit non-linear regression: *y =* -*4.9842 + 0.7268x*, *r^2^= 0.8692.* **B. Top, left:** cell-attached recordings from a PnO neuron after replacing standard ACSF with 9 mM KCl-ACSF (see *Methods*). Inset: experimental protocol. **Top, right:** example extracellular spike from the same recording, with individual raw traces (gray) superimposed and the average trace shown in black. Bottom: raster plot of 10 consecutive trials, aligned at t=-6.05 s before 1 mM GABA injection (arrowheads mark injection times; shaded area corresponds to the raw trace above). **C. Left**: whole-cell recordings of IPSCs evoked by paired-pulse stimulation in a representative experiment. Raw data from 20 trials are shown in gray, with their average in black. Inset: stimulation protocol. Oblique arrows indicate traces representing failures (i.e., no responses). **Right:** Scatter plot of R_1_ peak amplitudes and paired-pulse ratios (PPRs) from 51 observations. Bars are the mean ± SD **D.** Plot of PPR versus 1/CV² for R_1_. Non-linear regression yielded the best-fit curve described by: y = 0.8939 + 0.7136/x (N = 51, R² = 0.5271).

To pharmacologically isolate the GABA-mediated inhibitory post-synaptic currents (IPSCs), ionotropic glutamate receptors were blocked by adding kynurenic acid (5 mM, Sigma-Aldrich) to the perfusate. To improve the signal-to-noise ratio of chloride-mediated IPSCs, a ‘patch solution’ containing 150 mM KCl was used. The Nernst potential for Cl^-^ (E_Cl_) and hence, the reversal potential of the GABAergic IPSCs, was estimated as ∼ + 5.1 mV at 25°C. Series resistance (typically 10–20 MΩ) was estimated using −10 mV, 20 ms steps from the holding potential (Hamill et al., 1981); recordings were rejected when the series resistance shifted ≥20%. In cell-attached experiments, recording pipettes were filled either with a 150 mM K^+^-gluconate based solution - instead of KCl-, or with standard ACSF. Voltage-clamp mode was used in all cell-attached recordings; electrode command potential was 0 mV. Spontaneously firing neurons were recorded while perfusing the slice with regular ACSF (N=3) or with 9 mM KCl-ACSF (see Bracci et al. 1998), giving rise to a steady depolarization-evoked firing (2.2 ± 0.1 Hz, N=3, see **Fig. 1B**). Monopolar stimulating electrodes were pulled from borosilicate glass (tip diameter ≈5 µm), filled with 0.9% saline (NaCl), and positioned in the topographically defined LDT/PPT area—immediately below and near the medial pole of the superior cerebellar peduncle toward the midline and approximately 1 mm dorsal to the PnO core. Monopolar cathodic current pulses were delivered using a constant current unit (ISO-Flex, A.M.P.I.) connected to an Ag/AgCl wire inserted into the saline-filled glass micropipette. Postsynaptic responses were evoked using single pulses, paired-pulse stimulation (ISI=150 ms), or a short train of five pulses at ∼15 Hz, with stimuli delivered every 10 s. Stimulation parameters were controlled with pClamp 8.0 software (typical current pulses ranged from 10–500 pA with durations of 300–400 µs).

### Juxtacellular application of drugs

While recording PnO neurons, single doses of the following drugs were juxtacellularly injected by pressure application (typically 50-150 ms, 5-20 psi), through a glass ejection pipette connected to a Picospritzer III (Parker Instrumentation) and placed between 50 and 100 μm from the recording pipette: (±)-muscarine chloride hydrate (10 µM), (-)-nicotine hydrogen tartrate salt (20 µM), carbamoylcholine chloride (Carbachol, 10 µM), γ-aminobutyric-acid (GABA, 1 mM), 5- (aminomethyl)-1,2-oxazol-3-one (Muscimol, 100 µM), (±)-β-(Aminomethyl)-4- chlorobenzenepropanoic acid (Baclofen, 500 µM). Drugs were purchased from Sigma-Aldrich. Stock solutions were prepared in distilled water, stored at -8°C and diluted in 0,9% NaCl to their final concentration the day of the experiment.

### Analysis of IPSCs

Data were analyzed using Clampfit 10.7 (Molecular Devices) and Sigma Plot 11.0 (Systat Software Inc.). To analyze exclusively monosynaptic responses, our study included only IPSCs recorded in the presence of kynurenic acid (5 mM) that met, in addition, the following criteria (adapted from Rose and Metherate, 2005): i) low IPSC onset latency variability (typically SD < 2 ms); ii) no change in IPSC latency with increased stimulus intensity and iii) stable IPSC latency during repetitive (15 Hz) presynaptic stimulation. For a given recorded neuron, the mean peak amplitude of the first (R_1_) and subsequent evoked IPSCs (R_2_…R_5_) elicited by either single or multiple-stimuli protocols (i.e. paired-pulse or short trains), was calculated from 20 consecutive trials (at 0.1 Hz), by averaging the peak values of the corresponding IPSCs. Noise level was taken as ± 5 pA; lower amplitude peak events were considered failures. Given that synaptic failures were included in averaging, the mean peak amplitude of R_i_ (*i = 1…5*) strictly corresponds to the ‘synaptic efficacy’ of the *i-th* synaptic event. The latency, coefficient of variation (CV) and the time-constant (τ_off_) of the decay phase of R_1_ were routinely estimated; τ_off_ was estimated by fitting a single exponential function (y = a*e^-t/τ_off_) to the (90–10%) decay phase of the averaged IPSC. The activity-dependent short-term synaptic plasticity profile (short-term plasticity, STP) was evaluated by calculating the paired-pulse-ratio (PPR = R_2_/R_1_) from averages of 20 sweeps and the STP-ratio (STPR) in average responses to 20 successive applied trains; the STPR was calculated as a fraction of the mean value of R_3_, R_4_ and R_5_ relative to R_1_, i.e., STPR= [1/3 (R_3_+ R_4_+ R_5_)] / R_1_. The charge transfer (Q_inh_, pA.ms), a parameter of synaptic efficacy which incorporates the peak amplitude as well as the time course of synaptic events (Gardner, 1980; Hessler et al., 1993; Rinzel and Rall, 1974), was calculated as the time integral of the membrane current trace for each trial, excluding the stimulus artifact; the mean Q_inh_ was obtained from averages of 20 consecutive trials. Analysis of the effects of cholinergic agonists on IPSCs included: (i) the estimation of changes in synaptic efficacy by comparing evoked changes in functional synaptic parameters measured before (control) and after drug application during its maximal effect, (ii) the evaluation of the pre- or postsynaptic origin of changes in synaptic strength by quantifying changes in the PPR and the STPR, parameters representative of presynaptic function (Bekkers and Stevens, 1990; Davies et al., 1990; Débanne et al., 1996; Dobrunz and Stevens, 1997; Hess et al., 1987; Manabe et al., 1993; Metherate and Ashe, 1994; Schulz et al., 1994; Zucker, 1989, 1973) and changes in 1/CV^2^(R_1_) (Débanne et al., 1996; Faber and Korn, 1991; Malinow and Tsien, 1990). Spontaneous IPSCs (sIPSCs) were analyzed using Clampfit routines. The detection threshold of events was set at 5 pA. The cumulative probability distributions of amplitudes were constructed using 60–120 s gap-free recordings obtained before, immediately following and after (15 min) juxtacellular BACLOFEN application.

### Statistical Analysis

Statistical analyses were performed using SigmaPlot 11.0. The significance level was set at p < 0.05. Unless otherwise stated, data are presented as mean ± SE and shown in the figures as individual values (i.e., the average for each recorded neuron), where N represents the number of recorded neurons in each experimental series. Changes in synaptic efficacy were evaluated using either the paired Student’s t-test or the Wilcoxon signed-rank test, as appropriate, while unpaired data were analyzed using the unpaired Student’s t-test or the Mann-Whitney rank sum test.

## RESULTS

### IPSCs evoked by focal stimulation of the LDT/PPT area

Evoked IPSCs (eIPSCs) were elicited by repetitive stimulation of the region of brainstem tegmental nuclei and monitored via whole-cell, voltage-clamp recordings from PnO neurons held at -70 mV. The first eIPSCs (R_1_) from either stimulation pattern had a mean latency of 8.5 ± 0.3 ms, a mean amplitude (R_1(avg)_) of 76.8 ± 82.6 pA and a decay time constant of 24.8 ± 1.5 ms (N=51). eIPSCs exhibited a reversal potential of 8.5 ± 5.2 mV (N=3; **Fig.1A**), which closely matched the Nernst potential for Cl^-^ according to the composition of micropipette internal solution (see Methods). In addition, they were completely blocked by 100 µM picrotoxin (N=3, data not shown), further indicating mediation via GABA_A_-receptors. Under conditions preserving the physiological chloride concentration gradient (cell-attached recordings), juxta-cellular injections of 1 mM GABA transiently suppressed spontaneous or depolarization-induced action potential firing for approximately 2 seconds (**Fig. 1B**), indicating that GABA-mediated postsynaptic effects were inhibitory. In all experiments, eIPSCs were recorded after blockade of ionotropic glutamatergic transmission (5 mM kynurenic acid), a condition consistent with their monosynaptic nature.

### Short term plasticity profile of activated GABAergic synapses

Short-term use-dependent plasticity is a fundamental characteristic of synaptic connections, critically influencing neural network functionality. The dynamics of activated GABAergic synapses were assessed using two distinct stimulation protocols: paired pulses and brief short trains (5 pulses at 15 Hz; see Methods). In most neurons (40/51, ∼78%), facilitatory processes predominated (paired-pulse facilitation), with a paired-pulse ratio (PPR) ranging from 1.0 to 2.1, and a mean of 1.4 (± 0.2). Paired-pulse depression was observed in the remaining 11 neurons, with a PPR of 0.88 (± 0.10; range: 0.69–0.98). Overall, the mean PPR across all recorded neurons was 1.3 ± 0.3 (N = 51, **Fig. 1C**). Consistent with established models of use-dependent synaptic plasticity, paired-pulse facilitation -the predominant form of paired-pulse plasticity observed in our experiments-primarily reflects a low initial probability of neurotransmitter release (P_o_) (Débanne et al., 1996; Dittman and Regehr, 1998; Silver et al., 1998; Regehr, 2012). In line with this, the PPR exhibited an inverse correlation with 1/CV²(R1), an experimental proxy for P_o_ (McLachlan, 1978; Faber and Korn, 1991; **Fig. 1D**; *Pearson product moment* correlation: N = 51, r = -0.551, p <0.001). Additionally, also consistent with a low P_o_, R_1_ displayed a failure probability of 0.21 ± 0.03, together with substantial amplitude variability (CV(R_1_) = 48.1 ± 21.5%; see raw data in **Figs. 1C** and **2A**). Activation of GABA_B_ autoreceptors by GABA released in response to a presynaptic action potential acts as negative feedback on subsequent GABA release modulating the presynaptic terminal dynamics, thereby influencing paired-pulse plasticity (Davies and Collingridge, 1993; Kobayashi et al., 2012). This possibility was ruled out demonstrating that juxtacellularly-applied microvolumes of 100 µM baclofen, a GABA_B_-receptor agonist on eIPSCs, had no effect on R_1(avg)_ or the PPR (**Fig. S1A**; *paired t-test: N=9; R_1(avg)_: p=0.89; PPR: p=0.56*). However, baclofen was able to reduce the frequency of spontaneous IPSCs, indicating that GABA_B_ receptors may regulate additional inhibitory GABAergic inputs onto the same postsynaptic cells.

**Figure 2.**
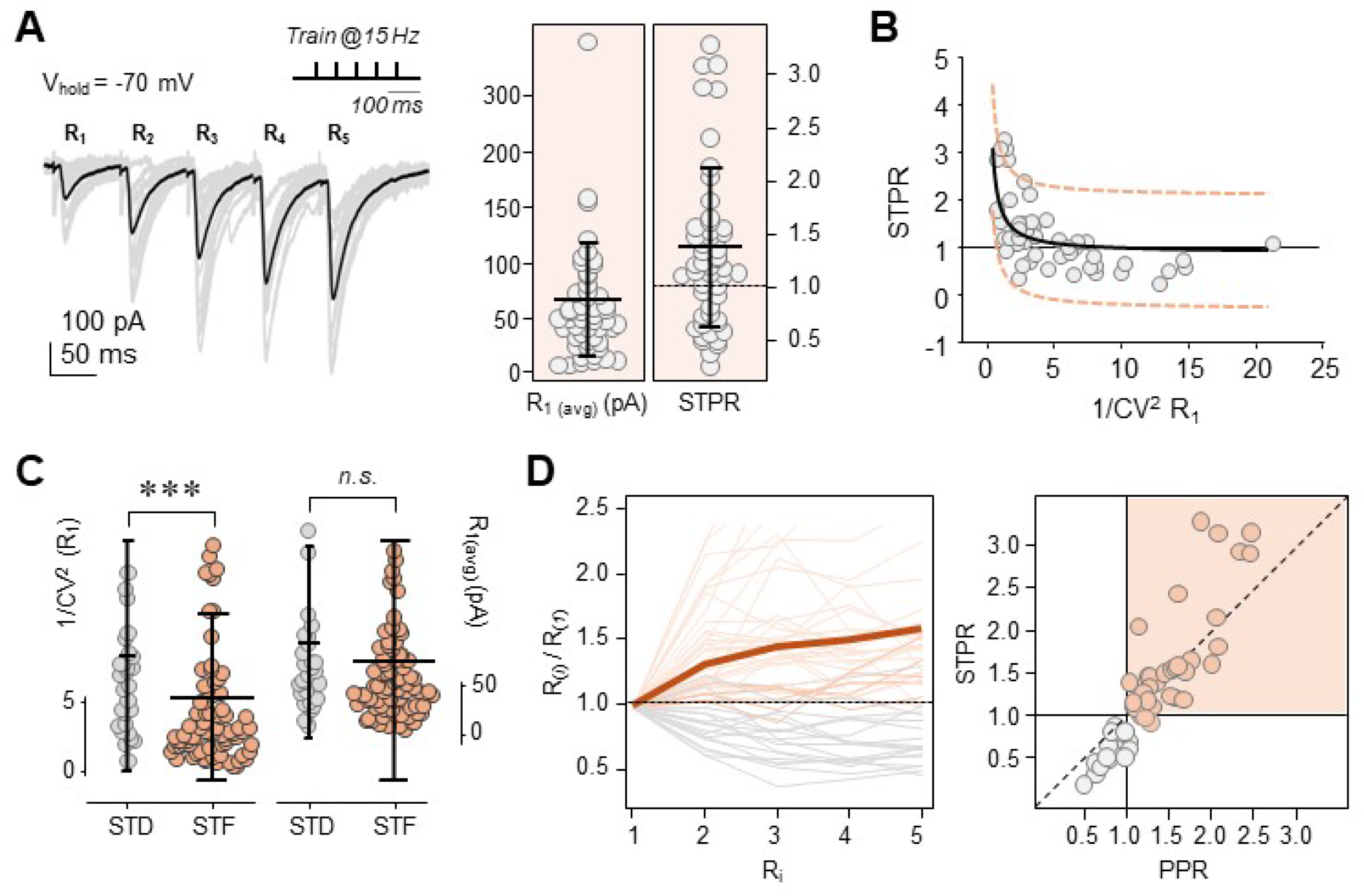
Short-term plasticity profile of activated GABAergic synapses during short-train stimulation. **A. Left**: representative whole-cell recordings of IPSCs evoked by a train of 5-pulses/15 Hz delivered in the LDT/PPT. Gray traces show 20 individual trials, and the black trace depicts their average. Inset: stimulation protocol. **Right:** Scatter plot of the mean peak amplitude of R_1_ (R_1(avg)_) and short-term plasticity ratios (STPRs). Each dot represents the mean value for a single experiment, while the bars indicate the mean ± SD. **B.** Plot of STPR versus 1/CV² for R_1_. A non-linear regression best-fit yielded the equation: y = 0.8679 + 1.0059/x (N = 57, R² = 0.4032). Discontinuous lines mark the 95% prediction band. **C.** Scatter plot of 1/CV² (R_1_) and of R_1(avg)_ pooled from both paired-pulse and short-train stimulation experiments (N = 108). Data are sorted according to the paired-pulse ratio (PPR) into two groups: short-term depression (STD; PPR < 1) and short-term facilitation (STF; PPR > 1). Bars depict the mean ± SD. *Mann-Whitney rank sum test: STD: N=29; STF: N=79; ***p<0.001; n.s.: not significant*. **D. Left:** Normalized peak amplitudes for pulses R1–R5 (values normalized to R1). The thick line represents the average profile. **Right:** Plot of STPR versus PPR for all experiments (N = 57); the shaded area indicates the region corresponding to short-term facilitation (STF; PPR > 1 and STPR > 1).

The spiking activity of GABAergic neurons in mesopontine areas -including the vlPAG and the LDT/SLD nuclei-varies across the wake-sleep cycle in a state-dependent manner (Boucetta et al., 2014; Weber et al., 2018). In naturally sleeping-waking rats glutamic acid decarboxylase-positive (GAD^+)^ neurons in the SLD exhibit REM sleep-specific (REM-S) and wake/REM sleep-specific (W/REM-S) maximal activity, firing in short bursts of action potential at ∼15–20 Hz (Boucetta et al., 2014). Therefore, short-term, use-dependent plasticity of activated GABAergic synapses was also examined using short trains stimulation (**Fig. 2A**), as an experimental proxy for physiological presynaptic activation (N=57). On average, eIPSCs exhibited frequency facilitation, with a short-term plasticity ratio (STPR, i.e.: [1/3 (R_3_+ R_4_+ R_5_)] / R_1_, see Methods) of 1.3 ± 0.7 (**Fig. 2A**, right). This average behavior -also illustrated in **Fig. 2D**-comprises a subset of neurons (36/57) displaying facilitation across the stimulation train (see the representative example **in Fig. 2A**, left) with a STPR of 1.7 ± 0.7 (range: 1.0 - 3.3). The remaining neurons (21/57) exhibited frequency depression with a STPR of 0.7 ± 0.2 (range: 0.3-1.0). As for the PPR, the STPR correlated negatively with 1/CV^2^(R_1_) (*Pearson product moment correlation: N=57, r=-0.525, p<0.0001,* **Fig. 2B**). Neither R_1(avg)_ (68.9 ± 53.0 pA, **Fig.2A**, right) nor PPR (1.2 ± 0.5, not shown) from this dataset differed significantly from paired-pulse stimulation datasets (*Mann-Whitney rank sum test: R_1(avg)_*: *p=0.744; PPR: p=0.225*). Therefore, data from paired-pulse and short train stimulation experiments (N=108) were pooled for further analysis of the presynaptic dynamics of activated GABAergic inputs. As expected, PPR and 1/CV^2^(R_1_) showed significant correlation (*Pearson product moment correlation: r*=-*0.257, p<0.01*). In contrast, the PPR and R_1(avg)_ were not correlated (*Pearson product moment correlation: r*=-*0.176, p=0.0681*) suggesting that short-term plasticity profiles are independent of R_1_ amplitude and thus unaffected by eventual variations of focal stimulation (Kim and Alger, 2001). This notion finds further support in the fact that neurons exhibiting paired-pulse facilitation (PPR>1) and paired-pulse depression (PPR<1) differed significantly in 1/CV^2^(R_1_) but not in R_1(avg)_ (**Fig.2C**; *Mann-Whitney rank sum test: PPD: N=29; PPF: N=79; 1/CV^2^(R_1_): p=0.001; R_1(avg)_: p= 0.084*). **Figure 2D** summarizes the short-term plasticity profiles of GABAergic synapses activated during the application of short trains stimulation protocols, highlighting the predominant short-term facilitation (i.e. PPR>1 and STPR>1, shaded area).

Collectively, these findings suggest that most GABAergic inputs activated by focal stimulation of the LDT/PPT area are mediated by a specialized subset of synapses characterized by a low release probability (as indicated by low 1/CV^2^ values for R_1_) and use-dependent facilitation (with PPR > 1 and STPR > 1) as the predominant short-term plasticity profile.

### Cholinergic modulation of GABAergic transmission at the PnO

In a first series of experiments (N = 28), we tested the effects of juxtacellular injections of the non-hydrolyzable ACh agonist carbachol (CCh, 1 mM) on GABAergic transmission (Fig. 3). Although CCh induced R_1(avg)_ changes in all 28 experiments, these effects were not uniform. In 20 out of 28 neurons, CCh reversibly depressed the eIPSC reducing R_1(avg)_ from 55.0 ± 33.4 to 37.9 ± 26.1 pA (**Fig. 3A1**). In the remaining 8 neurons, R_1(avg)_ increased from 104.6 ± 102.6 to 122.1 ± 117.08 pA (**Fig. 3B1**). These changes in synaptic strength were maximal near 9 min (9.1 ± 0.6 min) after juxtacellular CCh application and vanished (almost completely) by 20 min (19.2 ± 0.9 min).

**Figure 3.**
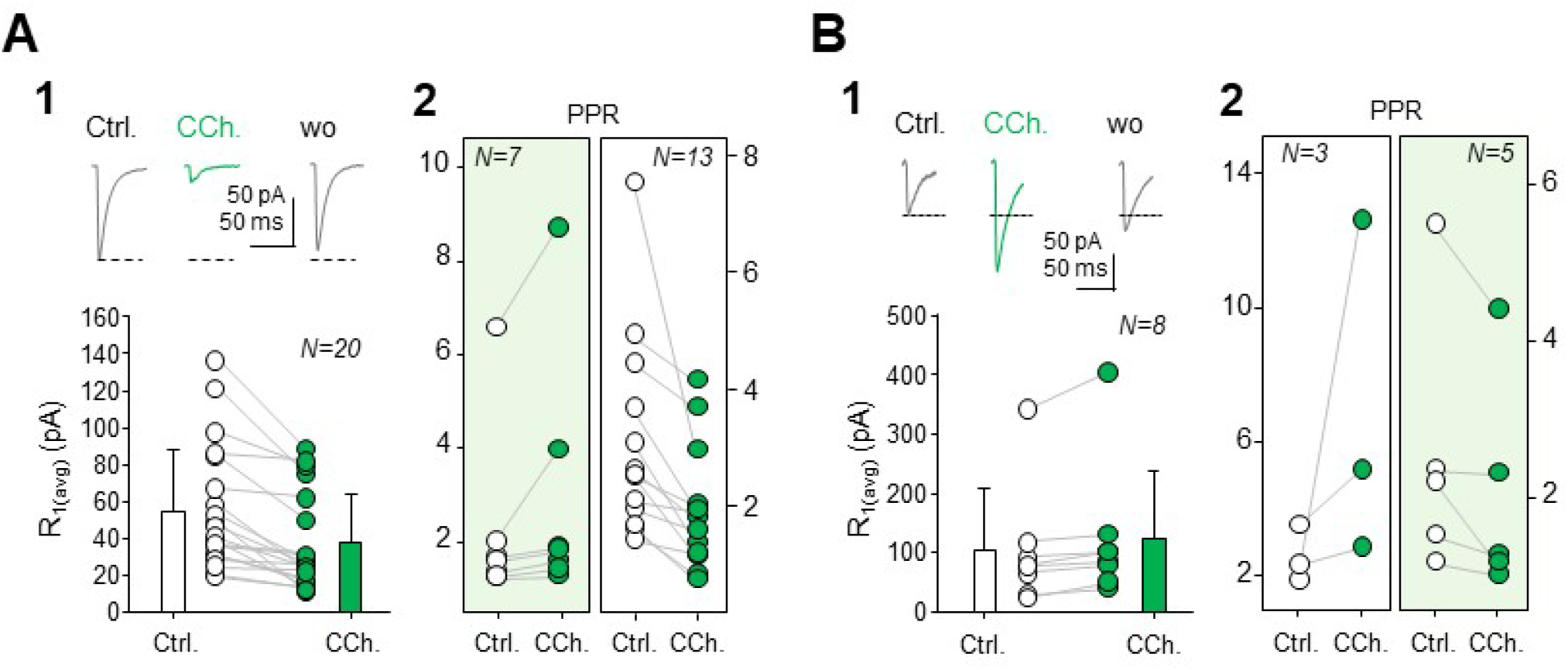
Modulation of activated GABAergic synapses by *carbachol* (CCh). **A. Synaptic depression. A1. Top**: whole-cell recordings of eIPSCs obtained before (control, Ctrl.) and after the juxtacellular injection of 1 mM carbachol (CCh.); the washout is also shown (wo). **Bottom:** Dot and Bar plot of R_1(avg)_ before (Ctrl.) and following CCh microinjections. Bars represent the mean ± SD. **A2**. Dot plot of PPR values sorted according to whether CCh induced an increase or a decrease relative to control. **B. Synaptic facilitation.** Same as A but illustrating CCh-induced increases of R_1(avg)_ (**B1**) and the corresponding changes in the PPR (**B2**). Sample size (N) is indicated for each graph. Shaded areas in A2 and B2 highlight the effects of CCh on PPR, consistent with presynaptic inhibition (A2) and presynaptic facilitation (B2); see main text for details.

To assess the possible presynaptic origin of CCh effects, we analyzed the PPR associated with changes in R_1(avg)_. In all experiments (28/28), modifications in synaptic strength were accompanied by changes in PPR, suggesting a presynaptic mechanism. However, both depression and facilitation were associated with either increases or decreases in PPR (**Fig. 3A2 and B2**). The expected increase in PPR indicative of a presynaptic locus of a CCh-mediated depression of transmission was observed only in 7 cells, in which the PPR increased from 1.21 ± 0.38 to 1.62 ± 0.61 (**Fig. 3A2, left**). In contrast, the remaining 13 neurons exhibited a decrease in PPR from 1.43 ± 0.35 to 1.22 ± 0.32 (**Fig. 3A2, right**). Similarly, facilitation of GABAergic transmission was accompanied by diverse changes in the PPR. In 5 neurons, the PPR decreased from 2.53 ± 1.72 to 2.02 ± 1.42 (**Fig. 3B2, right**), whereas in the remaining 3 neurons, PPR increased markedly from 2.51 ± 0.85 to 6.82 ± 5.2 (**Fig. 3B2, left**).

Taken together, the results described above suggest that CCh modulates the efficacy of activated GABAergic synapses via a complex presynaptic mechanism. Supporting this, the absence of CCh- mediated changes on the amplitude (87.2 ± 9.1% of control, N=10, p=0.19) and decay time constant (115.8 ± 7.6% of control, N=10, p=0.14) of postsynaptic currents evoked by juxtacellular 1 mM GABA injections (not shown) further supports the exclusion of any significant postsynaptic contribution to the CCh-induced modulation of GABAergic transmission in the PnO.

Activation of presynaptic muscarinic (mAChRs) and nicotinic (nAChRs) acetylcholine receptors can differentially modulate the initial release probability (P_₀_) of synaptic terminals, thereby influencing their short-term synaptic plasticity profiles. We hypothesized that the diverse effects of CCh on GABAergic transmission are mediated by the differential activation of presynaptic mAChRs and nAChRs. Accordingly, we evaluated the effects of selectively activating these receptors on evoked inhibitory postsynaptic currents (eIPSCs). Juxta-cellular application of muscarine (10 µM) reversibly decreased R_1(avg)_ in all experiments by 36.3 ± 18.9% (*p < 0.001, N=15;* **Fig. 4A1**), whereas nicotine (20 µM) produced an opposite effect, increasing R_1(avg)_ by 39.1 ± 24.7% (*p<0.001, N=14;* **Fig. 4B1)**. The maximal changes in R_1(avg)_ were observed at 7.6 ± 3.9 for muscarine and 2.8 ± 2.1 minutes for nicotine. Muscarine-induced decreases in R_1(avg)_ were consistently accompanied by increases in the PPR by 21.8 ± 5.1 % (*p=0.004; N=8*; **Fig. 4A2, left**) and the STPR by 93.3 ± 62.5 % (*p = 0.031, N=7*; **Fig. 4A2**, **middle panel**). In contrast, nicotine systematically reduced the PPR by 11.9 ± 1.9% (*p=0.003; N=8;* **Fig. 4B2, left**) and the STPR by 31.5 ± 2.2 % (*p = 0.031, N=6;* **Fig. 4B2, middle panel**). These effects did not correlate with the initial PPR or STPR values (see initial values in **Figs. 4A2** and **B2**).

**Figure 4.**
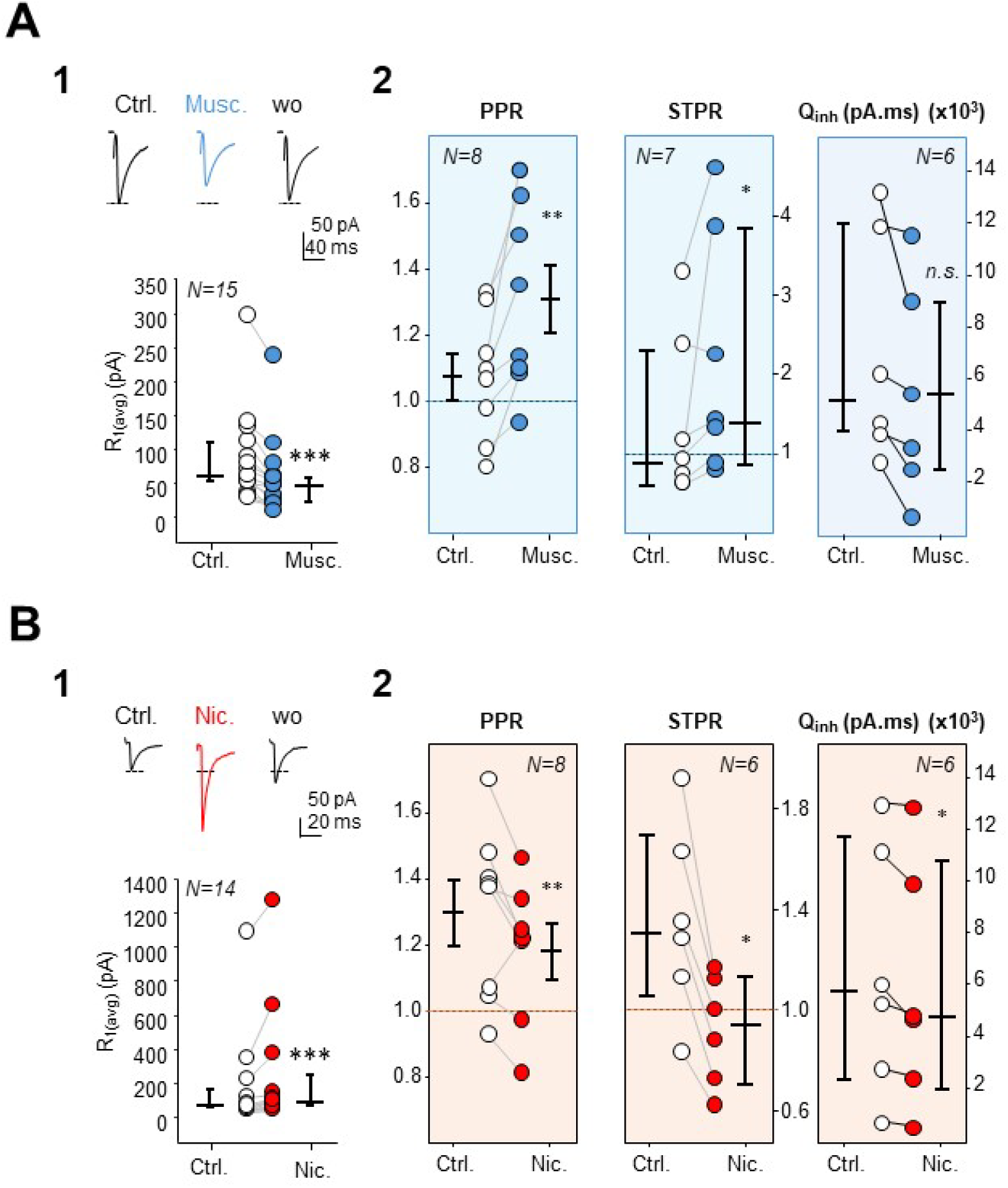
Modulation of activated GABAergic synapses by *muscarine* and *nicotine*. **A.** Modulation of synaptic efficacy and short-term plasticity by muscarine. **A1**. Top: Representative eIPSCs recorded before (Ctrl.) and immediately following juxtacellular injection of muscarine (Musc., 50-150 ms/20-30 psi, 10 µM); wo: washout. Bottom: R_1(avg)_ before (Ctrl.) and following the muscarine injection (Musc). **A2.** PPRs (left), STPRs (middle) and Q_inh_ (right) before (Ctrl.) and after muscarine injection (Musc.). **B.** Same as A but showing effects of nicotine (50-150 ms/20-30 psi, 20 µM). R_1_, PPR, STPR: *Wilcoxon signed-rank test, *** p<0.01;* Q_inh_: *paired t-test: * p<0.05, ** p<0.01; n.s: not significant;* Sample size (N) is indicated for each graph. Bars indicate the median ± the interquartile range.

The above results demonstrate that cholinergic modulation of activated GABAergic inputs to PnO neurons occurs via presynaptic mAChRs and nAChRs. Activation of these receptors elicited opposite changes in R1_(avg)_ and in the PPR/STPR, thereby changing presynaptic terminal dynamics. We next sought to determine whether muscarinic and nicotinic modulation leads to net depression or facilitation of the inhibitory effect on postsynaptic neurons during repetitive activation of GABAergic inputs. To this end, we quantified changes in the total inhibitory charge transfer (Q_inh_.) induced by muscarine (N = 6) and nicotine (N = 6) during short-train stimulation. The total charge transfer, which represents the cumulative movement of charge across the membrane during postsynaptic currents, serves as a proxy for the amount of neurotransmitter released, thereby providing an estimate of the net functional impact of synaptic inputs on postsynaptic outputs. Despite the opposing effects observed on R_1(avg)_, PPR, and STPR, both muscarine and nicotine consistently decreased Q_inh_, although the reduction observed with muscarine did not reach statistical significance (muscarine: 15.2 ± 18.7 %, *p=0.145, N=6*; nicotine: 13.7 ± 3.9 %; *p=0.038, N=6;* **Figs. 4A2 and B2, right panels**).

## DISCUSSION

In this study, we investigated whether and how cholinergic signaling modulates GABAergic synaptic input to REM-on neurons in the PnO, thereby contributing to the fine-tuning of REM sleep initiation, maintenance, and termination. Our main observation were: (1) focal stimulation in a restricted mesopontine area near the medial pole of the superior cerebellar peduncle evoked monosynaptic IPSCs at PnO neurons, likely mediated by GABA_A_ receptors, (2) the vast majority of these inputs (∼70%) result from activation of GABAergic terminals with low probability of transmitter release, short-term facilitation and no presynaptic modulation via GABA_B_ receptors, (3) these GABAergic inputs to PnO neurons are apparently endowed with functional presynaptic mAChRs and nAChRs whose activation modulates the probability of GABA release in opposite directions, with a muscarinic decrease and a nicotinic increase, and (4) for likely physiological presynaptic activity, however, activation of any receptor subtype leads to a net reduction of inhibition of PnO neurons.

### Characteristics of activated GABAergic inputs in putative REM-on PnO neurons

Previous electrophysiological studies of GABAergic actions on PnO neurons investigated membrane potential changes evoked by local application of GABA (Nuñez et al., 1998) and the modulation of spontaneous and proximally evoked IPSCs by carbachol (Heister et al., 2009). Weng et al. (2014) analyzed the cholinergic modulation of spontaneous postsynaptic currents but, specifically, in spinally projecting pontine reticular neurons. Here, we performed an in-depth analysis of functional properties and the cholinergic modulation of a subset of GABAergic inputs to putative REM-on PnO neurons. Our data indicates that recorded eIPSCs are caused by monosynaptic postsynaptic inhibition at PnO neurons, implying GABA_A_ receptors. The predominance of facilitation during paired-pulse and short train stimulation (**Figs. 1** and **2**), together with a relatively high CV(R_1_) and a considerable probability of failure of synaptic transmission, indicates that eIPSCs result from the activation of low-P_0_ GABAergic terminals (Bekkers and Stevens, 1990; Davies et al., 1990; Faber and Korn, 1991; Débanne et al., 1996; Dobrunz and Stevens, 1997).

Although the origin of the activated GABAergic projections was not specifically investigated in this study, the stimulation site suggests that both GABAergic neurons from the LDT and the axonal processes of GABAergic neurons of the vlPAG and the LPT were likely activated. Substantial evidence supports the role of these structures in controlling REM sleep executive areas (Semba, 1993; Sastre et al., 1996; Boissard et al., 2003; Lu et al., 2006; Vanini et al 2007; Sapin et al., 2009; Pino et al., 2017; Weber et al., 2018). Most GABAergic neurons in these regions are considered REM-off meaning their activity is drastically reduced during REM sleep and they are likely responsible for inhibiting REM sleep executive areas (Lu et al., 2006b; Sapin et al., 2009; Boucetta et al., 2014; Weber et al., 2018).

Among other physiological functions, short-term synaptic plasticity has been implicated in synaptic computation by filtering the transmission of information between neurons (Fortune and Rose, 2001). This property is particularly relevant during repetitive presynaptic activity. Synapses with a low initial probability of neurotransmitter release, exhibit frequency facilitation and function as high-pass filters, whereas synapses with a high initial probability of release, show frequency depression, acting as low-pass filters (Fortune and Rose, 2001; Abbott and Regehr, 2004; Monday et al., 2018). Here, the short-term plasticity profile of most GABAergic synaptic inputs to PnO neurons suggests that they may function as high-pass filters that favor GABAergic inhibition of neuronal targets during periods of repetitive presynaptic activity. Interestingly, evidence obtained *in vivo* indicates that the spiking activity of brainstem GABAergic cell groups thought to innervate REM sleep executive areas is strongly modulated by the sleep-wake cycle (Boucetta et al., 2014; Weber et al., 2018). Characteristically, REM sleep-related cells exhibit short periods of repetitive discharge of action potentials at 15-20 s^-1^ during REM sleep (Boucetta et al., 2014; Luppi et al., 2017; Jones, 2017; Weber et al. 2018; Blanco-Centurion et al; 2023).

### Presynaptic mAChRs and nAChRs mediate cholinergic modulation of activated GABAergic inputs

In the present study, we found that CCh consistently elicited transient changes in the efficacy of GABAergic synapses. Although this phenomenon was observed in all tested PnO neurons, the changes in R_1(avg)_ amplitude were not homogeneous: CCh reduced R_1(avg)_ amplitude (depression) in about two-thirds of neurons while enhancing R_1(avg)_ (facilitation) in the remaining cells. In all cases, changes in R_1(avg)_ were accompanied by changes in the PPR, indicative of a presynaptic mechanism. This conclusion is further supported by the absence of changes in membrane currents evoked by the juxtacellular application of GABA microvolumes in the presence of CCh. Our findings contrast with those of Heister et al. (2009), who reported no effect of CCh on evoked IPSCs. This discrepancy may stem from their stimulation site (50–200 µm from the recorded soma), which might have preferentially activated local PnO GABAergic interneurons with a probably distinct profile of cholinergic modulation. Altogether, our data provide further experimental support for a functional interaction between cholinergic and GABAergic innervation of the PnO, as reported in several previous studies (e.g., Sandfor et al., 2003; Xi et al., 2004; Marks et al., 2008; Vanini et al., 2011; reviewed in Watson et al., 2011 and Brown et al., 2012).

Due to its non-selectivity, CCh likely exerts a range of effects on GABAergic synaptic transmission, including those mediated by the activation of presynaptic muscarinic (mAChRs) and nicotinic (nAChRs) receptors -with muscarine replicating synaptic depression and nicotine facilitating transmission. However, the effects of CCh on the short-term plasticity profile associated with modulations in R_1(avg)_ challenge this interpretation. A net prevailing muscarinic or nicotinic presynaptic effect of CCh might account for results observed in 43% of neurons (opposite shifts in R_1(avg)_ and the PPR) whereas in the remaining cells, CCh effects on R_1(avg)_ and the PPR are compatible with a diverse combination of muscarinic or nicotinic presynaptic effects.

mAChRs are G protein-coupled receptors that can be subdivided into two groups (Assous, 2021). The M1-like receptors group (including M1, M3, and M5 receptors) are coupled to Gq/11 G-proteins, and usually associated with excitatory neural effects mediated by activation of protein kinase C and phospholipase C. The M2-like receptors group (including M2 and M4 type receptors) are coupled to Gi/o G proteins inhibiting adenylyl cyclase and are implicated in inhibitory effects. At synaptic level, activation of the M2-like group mediated presynaptic inhibition through inhibition of voltage-dependent Ca^2+^ channels leading to a reduction of evoked neurotransmitter release probability. Neuronal nAChRs are ionotropic receptors permeable to Na^+^, K^+^, and Ca^2+^ ions that mediate fast synaptic transmission and, most importantly, intracellular signaling via Ca^2+^- dependent signaling pathways (Wonnacott, 1997; Dajas-Bailador and Wonnacott, 2004; Gotti et al., 2009; Zoli et al., 2018). Activation of nAChRs at presynaptic level causes a rise in the cytosolic Ca^2+^ leading to an increase in release probability via the modulation of several critical steps in neurotransmitter release. At the brainstem mesopontine region, a large body of evidence indicates the existence of cholinoceptive structures whose pharmacological manipulation causes changes in the occurrence and timing of REM sleep episodes (reviewed in Brown et al., 2012; Luppi et al., 2012; Weber et al., 2018; Wang et al., 2021). Experimental data obtained using different approaches including *in vivo* and *in vitro* models, indicate that cholinergic modulation of sleep related mesopontine structures may involve both mAChRs (M1 and M2-like groups) and nAChRs (probably α4β2) (Greene et al., 1985; Stevens et al., 1993; Gillin et al., 1993; McCarley et al., 1995; Brischoux et al., 2008; Léna et al., 2004; Coleman et al., 2004; Heister et al., 2009; Brown et al., 2012).

Our data indicate that GABAergic terminals contacting PnO neurons are modulated by two distinct cholinergic receptor systems. Activation of M2-like mAChRs produces presynaptic inhibition, as evidenced by a reduction in R1_(avg)_ and corresponding increases in both PPR and STPR. In contrast, activation of nAChRs mediates presynaptic facilitation, as indicated by an increase in R_1(avg)_ coupled with reduced PPR and STPR. Given that mAChRs and nAChRs utilize independent intracellular signaling mechanisms, co-activation of both receptor types by CCh -when they are co-expressed at the same presynaptic terminals- may result in a combination of their respective cellular effects. Moreover, dual muscarinic/nicotinic effects of CCh -characterized by shifts in both R_1(avg)_ and the short-term plasticity profile in the same direction observed in 57% of neurons, may depend on the simultaneous activation of mAChRs and nAChRs at these terminals. Modulation of synaptic transmission mediated by activation of different presynaptic receptor subtypes with opposing effects on transmitter release has been described in other neural structures (Zhang and Warren, 2002; Bono et al, 2023). The predominance of the effects mediated by one receptor over the other on R_1(avg)_ and the short-term plasticity profile depends, among other factors, on each receptor type’s expression level and the functional state of the terminals (including intracellular signaling pathways; Perrins and Roberts, 1994; Grilli et al., 2009; Marchi and Grilli, 2010).

It is worth noting that despite the opposing effects of presynaptic activation of mAChRs (depression) and nAChRs (facilitation) on GABAergic transmission at the PnO, activation of either cholinergic receptor subtype during presynaptic activation mimicking physiological activity patterns of REM-off GABAergic neurons (Boucetta et al., 2014; Weber et al., 2018), reduced the inhibitory drive onto PnO neurons (as represented by Q_inh_ in **Fig. 4**). Thus, it is reasonable to speculate that an increased cholinergic tone during the transition from non-REM to REM sleep leads to a local inhibition of GABA release via the co-activation of presynaptic mAChRs and nAChRs. This scenario is plausible because volume transmission likely constitutes the dominant mode of transmission at the target regions of cholinergic projections in brainstem neural circuits involved in REM sleep control (Saper and Fuller, 2017), as well as in other CNS regions (Picciotto et al, 2012; Bennett et al., 2012; Lendvai and Vizi, 2008; Yamasaki et al., 2010; Liang and Marks, 2014). Although our data strongly suggest that mAChRs and nAChRs are co-expressed in the same GABAergic presynaptic terminals, a definitive demonstration requires further analysis of cholinergic modulation of GABA transmission using more selective stimulation, such as the minimal stimulation method, which activates a single or only a few presynaptic axons (Raastad et al., 1992).

### Possible functional implications of our data

The mutual inhibition model (or flip-flop switch model; Lu et al., 2006) has provided seminal conceptual frameworks for advancing the study of sleep mechanisms (Fuller et al., 2007; Brown et al., 2012; Weber and Dan, 2016; Héricé et al., 2019; Weber, 2017). This model consists of mutually inhibitory interactions between REM sleep-promoting and REM sleep-inhibiting neuronal structures. Although recent evidence points to dual GABAergic/glutamatergic control of REM sleep-promoting mesopontine structures, the REM sleep sleep-promoting effect of cholinergic innervation of these structures interacting with GABAergic REM sleep-suppressive influences still has considerable support (Mark and Sinton, 2010; Vanini et al., 2011; Arrigoni et al., 2016; Peever and Fuller, 2017; Weber et al., 2018; Arrigoni and Fuller, 2019; Grace and Horner, 2020). While the application of cholinergic agonists in this area is associated with a REM sleep facilitating effect (not triggering), GABA agonists have a clear REM sleep-suppressive effect (see *Introduction*). A local mutual inhibition between GABAergic and cholinergic inputs to the PnO is consistent with their antagonistic effect on REM sleep. Inhibition of ACh release from cholinergic terminals by GABA has been proposed based on electrophysiological, pharmacological, and immunolabeling evidence (Vazquez and Baghdoyan, 2004; Marks et al., 2008; Flint et al., 2010; Vanini et al., 2011; Liang and Marks, 2014). In this study, we demonstrate that GABAergic terminals in contact with PnO neurons are under intricate cholinergic presynaptic regulation, in line with a design of local, presynaptic mutual inhibition between GABAergic and cholinergic inputs to the PnO.

The transition from non-REM to REM sleep is characterized by a progressive increase in the activity of REM-on neurons in the mesopontine reticular region under cholinergic influence, along with a progressive reduction in the firing rate of REM-off GABAergic neurons of the vlPAG and the LPT, mediating the gating of REM sleep (Weber et al., 2018). The net presynaptic inhibition of GABAergic inputs mediated by the activation of cholinergic receptors may have a role in controlling the characteristics and duration of this transition (Fraigne and Peever, 2015). Moreover, together with the presynaptic GABAergic inhibition of cholinergic terminals at the PnO, the cholinergic relief of GABAergic inhibition could offer a mechanism of local mutual inhibition at the presynaptic level between GABAergic and cholinergic terminals at the PnO, with possible relevant functional consequences on the ultradian rhythm of non-REM/REM alternation (Brown et al., 2012; Luppi et al., 2012; Weber et al., 2018; Wang et al., 2021).

## Acknowledgements

Special thanks are due to Dr. Jack Yamuy, whose inspiration, creativity, dedication, and support were indispensable to the completion of this study. The authors also wish to thank Drs. Ángel Núñez and Verónica Egger for their valuable comments on the manuscript. This research was supported by FCE2009_3115-ANII, CSIC-CAP, UdelaR, and PEDECIBA.

## Author contribution

E.P., H.K. and M.B. conceived and designed research; E.P. and H.K. performed experiments; E.P., H.K. and M.B. analyzed data and interpreted results of experiments; E.P. and H.K. prepared figures. E.P. drafted the paper and E.P. and M.B. wrote, reviewed and edited the final version of the manuscript. All authors approved its final version.

---

**Figure S1.**
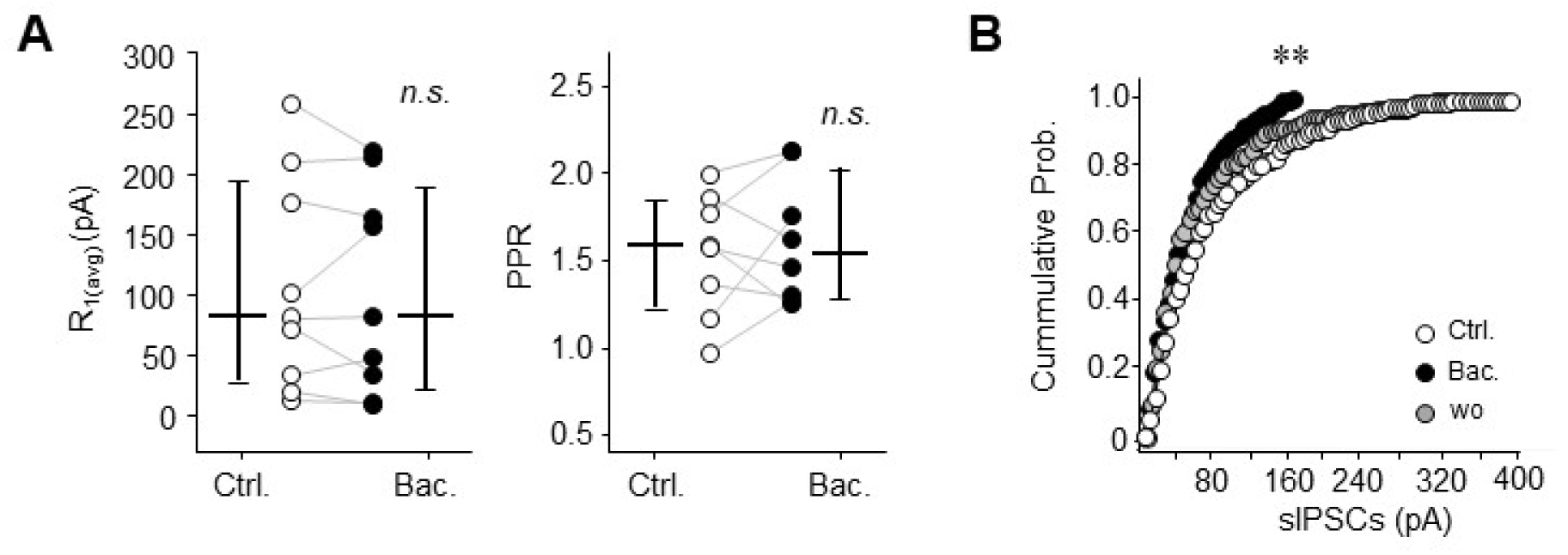
Effects of GABAB-receptor activation on GABAergic synaptic inputs to PnO neurons. **A.** R_1_ peak amplitudes (left) and PPRs (right) before (Ctrl.) and immediately following juxtacellular injection of 100 µM baclofen (50-150 ms/20-30 psi, Bac.); paired t-test: N=9, n.s.: not significant. *Bars are the median ± the interquartile interval*. **B**. Cumulative probability distribution of amplitude of sIPSCs sampled during a representative experiment illustrated in **A**; Plots were constructed by detecting consecutive events during 60-120 s gap-free recordings sampled before (Ctrl.), immediately following (Bac.) and 15 min after (wo: washout) baclofen injection. Immediately after Baclofen injection, the cumulative probability distribution shifted leftward compared to control (*Kolmogorov–Smirnov test: ** p<0.005*) indicative of a reduction in event amplitude with a suppression of larger events.

## Notes

### Competing Interest Statement

The authors have declared no competing interest.

### Summary of Updates

- English language editing and style refinement - Rewording of the description, statistical analysis, and graphical representation of carbachol's effects on GABAergic transmission (Figure 3) - Correction of inaccurate citations - Figure S1 revised.

